# Evolution in response to prophage activation attenuates the virulence of culturable *Serratia symbiotica* relatives of aphid endosymbionts

**DOI:** 10.1101/2024.12.04.626866

**Authors:** Anthony J. VanDieren, Jeffrey E. Barrick

**Affiliations:** Department of Molecular Biosciences, Center for Systems and Synthetic Biology, The University of Texas at Austin, Austin, Texas, USA

## Abstract

*Serratia symbiotica* bacteria exhibit a range of relationships with aphids. They may be co-obligate mutualists, commensals, or even pathogens depending on the strain, aphid host species, and environment. *Serratia symbiotica* CWBI-2.3^T^ (CWBI), a culturable member of this group, is transmitted to embryos transovarially when it is injected into pea aphids (*Acyrthosiphon pisum*), the same route used by *S. symbiotica* strains that are vertically inherited endosymbionts. Yet, aphids colonized with CWBI die before they give birth to infected offspring. We evolved laboratory populations of CWBI through 15-30 serial passages at two different temperatures in rich media. These nutrient-replete conditions mimic aspects of the environment within aphid hosts that lead to the evolution of reduced endosymbiont genomes. Unexpectedly, all *S. symbiotica* populations propagated at one temperature appeared to evolve slower growth after only a few days due to reactivation of a lytic prophage from the CWBI genome. Though these populations continued to reach saturating cell densities slower than cultures of the ancestor throughout the experiment, representative clones isolated from them had mutations affecting lipopolysaccharide biosynthesis and were resistant to the phage. Some evolved strains exhibited less virulence when injected into aphids, and we observed instances of gene inactivation and loss mediated by insertion elements. Our results illustrate how transposons and prophages can dominate laboratory evolution of newly cultured bacteria, particularly those that are host-associated in nature and have genomes rife with selfish DNA elements. They also suggest that bacteria-phage coevolution can catalyze evolutionary paths that contribute to converting pathogens into stably inherited endosymbionts.

**IMPORTANCE:** Laboratory experiments can be used to explore evolutionary innovations in how microbes associate with animal hosts. *Serratia symbiotica* bacteria exhibit a variety of interactions with aphids. Some strains are obligate endosymbionts. Others have facultative associations with benefits or costs depending on the environmental context. *S. symbiotica* CWBI-2.3^T^ (CWBI) resembles aphid endosymbionts in how it can be transovarially transmitted to aphid embryos. However, adults injected with CWBI do not survive long enough to give birth to infected offspring. We evolved this aphid protosymbiont in rich media to see if this would attenuate its virulence and recapitulate genome reduction observed in endosymbionts. We observed large deletions and gene inactivation, but reactivation of a prophage from the CWBI genome and then evolution of phage resistance dominated. Some evolved strains became less virulent to aphids, suggesting that evolution driven by selfish DNA elements can contribute to the emergence of new endosymbionts from pathogen ancestors.

## INTRODUCTION

Microbes have evolved a wide variety of associations with insects, including living as endosymbionts within the cells and tissues of their hosts (1–3). Insect symbionts can be broadly classified as obligate or facultative in terms of their relationships with their hosts. Obligate symbionts are required for host survival. They typically colonize specialized host cells or organs, are reliably vertically transmitted to offspring, and evolve severely reduced genomes that make it difficult or impossible to axenically culture them. Facultative symbionts are not required for host survival and can be vertically or horizontally transmitted. They often display complex relationships with their hosts. For example, a facultative symbiont may impose a general fitness cost on its host, but provide a benefit, such as protection against parasitoids, that can outweigh this cost under certain conditions (4). Facultative symbionts often have larger genomes than obligate symbionts, and some can be cultured outside of their insect hosts.

Pea aphids (*Acyrthosiphon pisum*) have both obligate and facultative endosymbionts. The obligate symbiont (*Buchnera aphidicola*) is required for aphid survival because it synthesizes essential amino acids that are not present in sufficient quantities in their diet (5, 6). *Buchnera* has a greatly reduced genome and has not been cultured. Facultative symbionts of pea aphids—including *Hamiltonella defensa, Regiella insecticola*, and *Serratia symbiotica*—colonize bacteriocytes and can also be found in the hemolymph and various tissues (6). Different strains of the bacterium *Serratia symbiotica* exhibit a range of relationships with aphids. Some *S. symbiotica*, like those found in the aphid *Cinara cedri*, are co-obligate symbionts with reduced genomes and characteristics similar to *Buchnera* (7–10). Others, like *S. symbiotica* Tucson in *A. pisum*, are facultative, providing a series of context dependent benefits, such as protection against heat stress and parasitoid wasps (4, 11, 12).

Recently, the first culturable strains of *S. symbiotica* have been isolated from aphids. These culturable *S. symbiotica*, which include strain CWBI-2.3T (CWBI), primarily colonize the aphid gut and may be horizontally transmitted between aphids feeding on the same plant (7, 13–16). They have genomes that are intermediate in size between those of other culturable *Serratia* species, such as *Serratia marcescens*, and uncultured facultative symbionts, like *S. symbiotica* Tucson (10, 17, 18). When injected into aphids, CWBI is transovarially transmitted to developing aphid embryos in the same manner as *Buchnera* and *S. symbiotica* Tucson (10). However, CWBI is unable to establish stable vertical transmission because infected adults do not survive long enough to give birth to infected offspring (10). Because of its latent capacity for transmission, CWBI and similar culturable strains like *S. symbiotica* HB1, may represent aphid protosymbionts that are poised to evolve into vertically inherited endosymbionts (10, 18, 19).

It has been hypothesized that endosymbionts arise from pathogenic ancestors and that evolving reduced virulence is necessary for transitioning to this new lifestyle (20–22). If we could evolve or engineer CWBI to be less virulent, this might complete its transformation into a vertically inherited aphid endosymbiont. Passaging microbes *in vitro* is a classical method for attenuating pathogens. Key virulence factors are often genetically unstable and rapidly lost in an environment in which they provide no benefits. Laboratory passaging has been used to develop live attenuated bacterial vaccines (23–26), and unintentional losses of virulence have occurred in pathogens when they were cultured in the laboratory (27–30). Mutations that reduce virulence may evolve due to random genetic drift if populations are subjected to single-cell bottlenecks as they are passaged, but this process is slow because bacterial mutation rates are low. Alternatively, serially passaging large populations of microbes can rapidly select for mutants with increased growth rates *in vitro* and possibly, thereby, reduce virulence *in vivo*. Both types of experiments can lead to chromosomal deletions and gene loss, potentially mimicking mechanisms of genome evolution experienced by transitional symbionts in nature.

We performed serial transfer evolution experiments with CWBI to examine what mutations accumulated in its genome during adaptation to laboratory culture conditions. We evolved populations at two different temperatures to observe more varied evolutionary trajectories. Reactivation of a prophage occurred in all populations in the lower temperature treatment, followed by the evolution of resistance to this phage through mutations in genes involved in lipopolysaccharide biosynthesis. Insertion sequence activity led to gene inactivation and loss in evolved strains, but not at as high of a rate as might be expected from their prevalence in the CWBI genome. Aphids survived for longer after they were injected with some strains evolved at the lower temperature compared to the ancestor. This attenuated virulence, if compounded by further evolution, might eventually lead to stable vertical transmission. Our results illustrate how mobile DNA elements, such as transposons and prophages, which may proliferate in the genomes of host-associated bacteria (17, 31, 32), shape how they evolve. They further suggest that coevolution with these elements may be important for the emergence of new endosymbionts from pathogen ancestors.

## RESULTS

### *Serratia symbiotica* laboratory evolution experiment

CWBI-GFP is a genetically modified variant of *Serratia symbiotica* CWBI-2.3^T^ with a cassette expressing GFP integrated into its chromosome (10). We evolved CWBI-GFP by serially propagating twenty-four populations through cycles of 1:1000 dilution and regrowth in a rich medium (tryptic soy broth) (**Fig. 1A**). Twelve of the populations (LT-01 through LT-12) were cultured at a lower temperature, the ambient temperature of 22–25°C in our laboratory. The other twelve (HT-01 through HT-12) were cultured at a higher temperature, in an incubator set to 30°C. The CWBI-GFP ancestor grew to saturation within 24 hours at both temperatures, and all populations were initially transferred daily. Two of the 30°C populations (HT-05 and HT-12) were eventually discontinued due to suspected contamination.

**Fig. 1.**
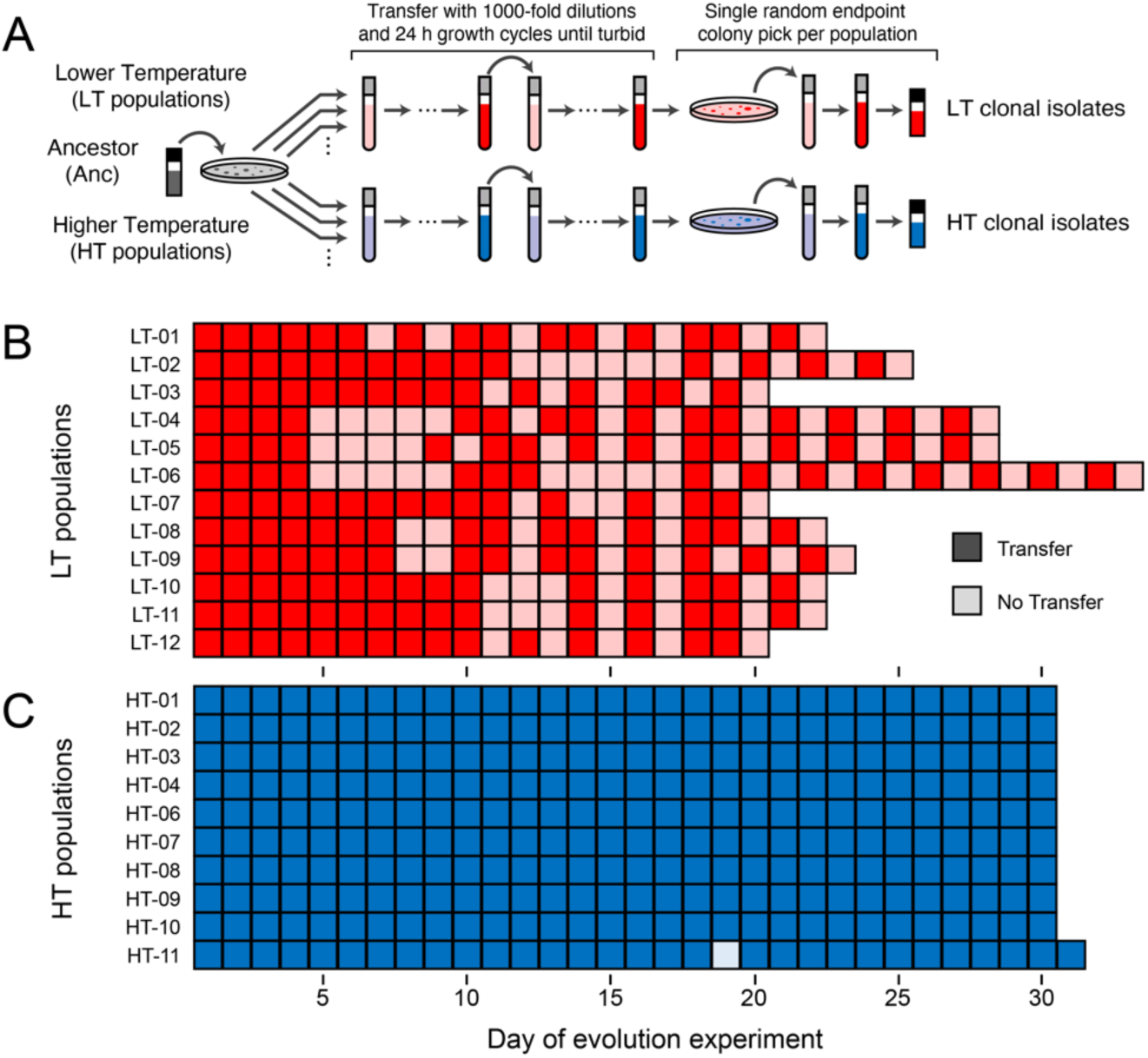
*Serratia symbiotica* CWBI-3.2^T^ evolution experiment. (**A**) Procedures used to begin the independent populations of the evolution experiment, carry out cycles of serial transfer and regrowth, and isolate endpoint clones for characterization. (**B**) Transfer histories of lower temperature (LT) populations evolved at ambient laboratory temperature (22–25°C). (**C**) Transfer histories of higher temperature (HT) populations evolved in an incubator at 30°C. In panels B and C, darker shading indicates that a population was transferred to fresh medium with a 1:1000 dilution on that day. Lighter shading is used for days when a population was not transferred because it had not yet reached the normal turbidity expected for a saturated culture of the ancestor strain.

### Delays in population growth always evolved in the LT treatment

For every LT population, we observed a pronounced delay in how long it took the cultures to reach normal saturating turbidity levels beginning between 4 and 10 passages (**Fig. 1B**). These delayed cultures were considerably less turbid than the CWBI-GFP ancestor after 24 hours and sometimes contained stringy debris resembling cobwebs. All eventually reached an apparently saturating turbidity after additional days of incubation, at which point they were transferred. After further variability in the delays observed on subsequent days, all LT populations began to consistently reach full turbidity between 24 and 48 hours of incubation. We propagated all LT populations for a total of 15 transfers, which took from 21 to 33 days.

Most HT populations continued to reach saturation within 24 hours for the duration of the evolution experiment and were propagated through a total of 30 transfers (**Fig. 1C**). The one exception was population HT-11, which exhibited a growth delay on day 19, but then returned to reaching saturation each day. On day 20 and thereafter, cells in this population rapidly settled to the bottom of the test tube when it was removed from the shaking incubator. To further investigate this new phenotype, we randomly selected ten colonies from the HT-11 population for characterization at the end of the experiment. Saturated cultures grown from all ten of these clonal isolates exhibited the same rapid settling as cultures of the full population.

### Growth rates of evolved LT populations and clonal isolates are poorly correlated

To characterize CWBI evolution, we picked a single clonal isolate from each endpoint population (**Fig. 1A**). We recorded growth curves for all populations and these clonal isolates in microwell plates using a plate reader (**Fig. 2**). Nine of the LT populations evolved at 22–25°C had significantly slower exponential growth rates than the ancestor at 25°C (**Fig. 2A**). The evolved LT clones exhibited more variability in their growth rates compared to their populations of origin (**Fig. 2B**). Some grew more rapidly, while others grew much more slowly than their populations of origin. The net effect was that, on average, population growth rates were not significantly different from clone growth rates (*p* = 0.283, ANOVA *F*-test), even though the growth rates of individual clones relatively poorly correlated with the growth rates measured for their populations of origin (*r*_xy_ = 0.465, sample Pearson’s correlation coefficient).

**Fig. 2.**
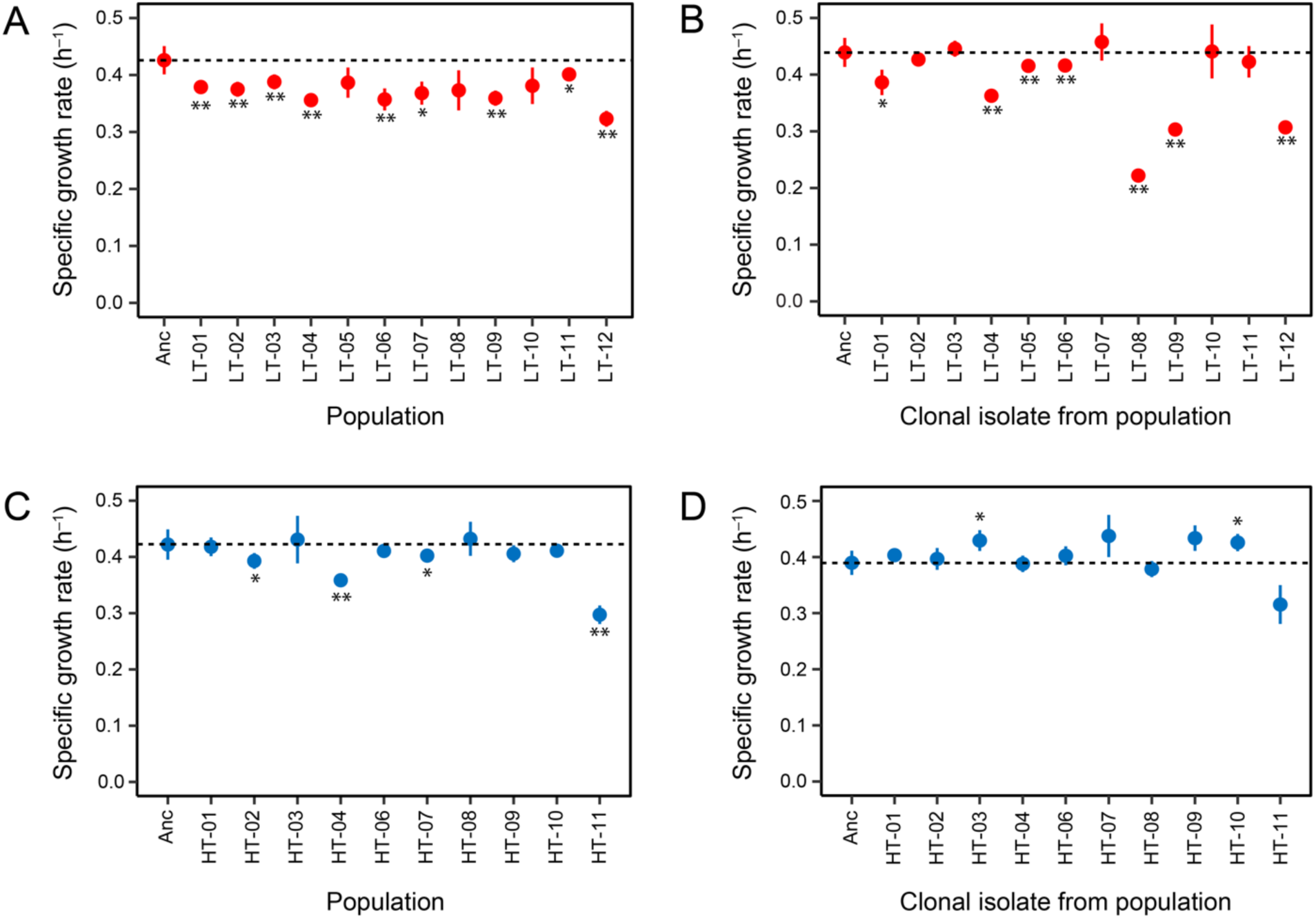
Growth rates of evolved populations and strains. Exponential growth rates estimated from microplate assays for (A) endpoint LT populations at ambient room temperature, (B) clonal isolates from the LT populations at ambient room temperature, (C) endpoint HT populations at 30°C, and (D) clonal isolates from the HT populations at 30°C. Horizontal dashed lines are reference lines at the growth rate of the CWBI-GFP ancestor strain (Anc). Error bars are standard deviations. Asterisks above or below points indicate that there is a significant difference between the measured growth rate of a population or clonal isolate and the ancestor in the same microplate assay (two-tailed *t*-tests with Bonferroni correction, \**p* < 0.05 and \*\**p* < 0.01).

There were fewer changes in the evolved samples and less variability between them in the 30°C treatment. A majority of the evolved HT population had growth rates that were indistinguishable from that of the ancestor strain, with just four exhibiting significantly reduced growth rates (**Fig. 2C**). Evolved HT clone growth rates (**Fig. 2D**) also correlated better with their populations of origin (*r*_xy_ = 0.766) than was the case for the LT samples. The main outlier was population HT-11 and its clonal isolate, which both appeared to grow more slowly, likely due to the rapid settling phenotype interfering with measuring growth from changes in culture turbidity. Though there was no difference, on average, in the growth rates of the HT clones and populations (*p* = 0.605, ANOVA *F*-test), the HT-03 and HT-10 clones individually exhibited significantly increased growth rates compared to the ancestor (**Fig. 2D**).

### Phages derived from a CWBI lysogen are present in evolved LT populations

The final LT populations took more than 24 h to reach saturation and accumulated stringy debris. We hypothesized that this debris was derived from cells lysed by phages, and that partial lysis of the cell populations was causing the apparent growth delays. When we spotted phage lysates that we made from LT endpoint populations on top agar containing the CWBI-GFP ancestor, we observed turbid zones of clearing after 48 h of incubation that supported this hypothesis (**Fig. 3A**). Transmission electron microscopy (TEM) imaging of lysates showed that they contained tailed phages (**Fig. 3B**). These phages have a capsid with a diameter of ∼70 nm and a tail with a length of ∼150 nm. The diameter of the tail is ∼10 nm except for a ∼50 nm segment near the end where it widens to ∼20 nm.

**Fig. 3.**
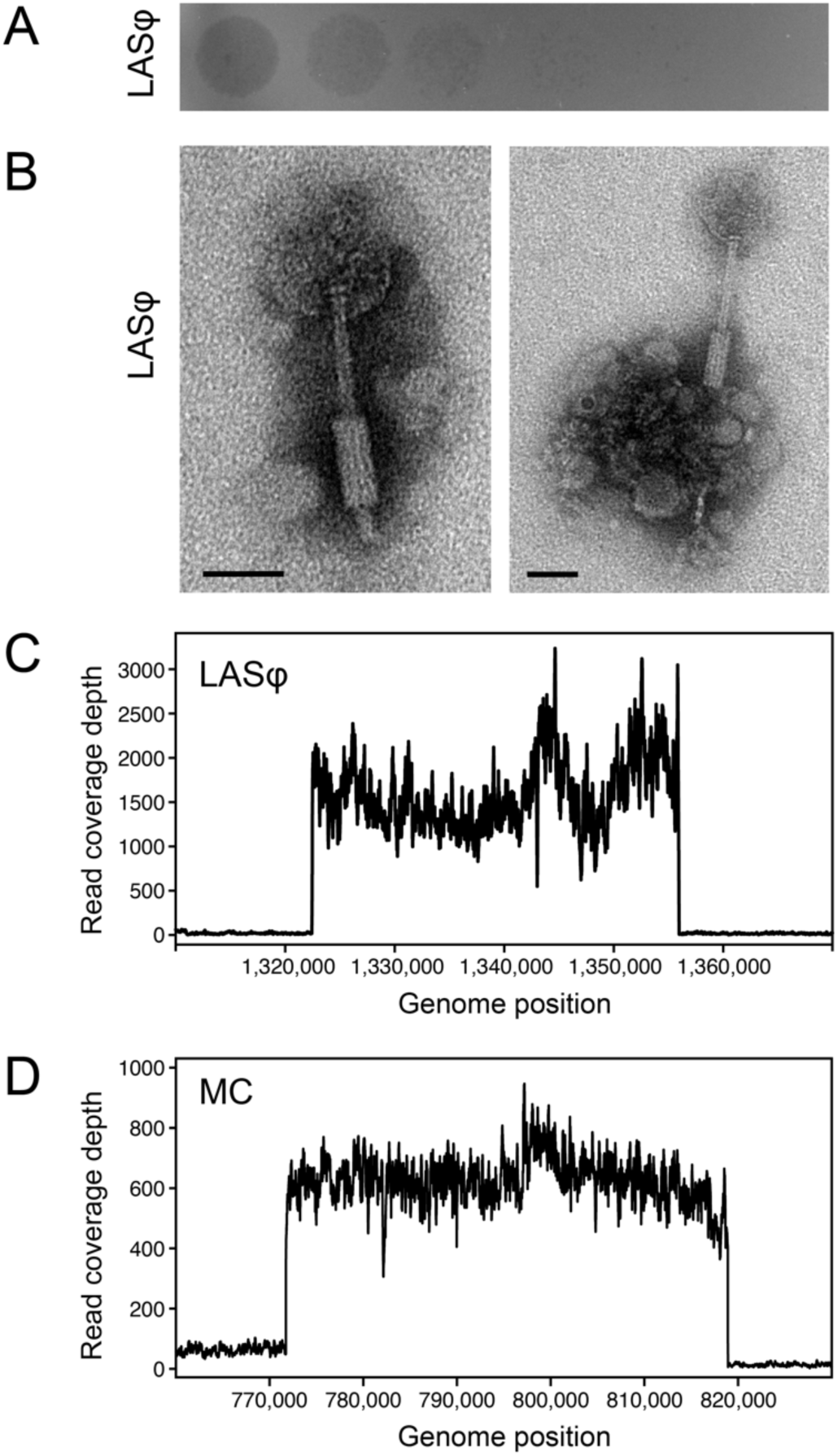
Activation of *S. symbiotica* CBWI prophages. (**A**) Turbid plaques formed by lysate containing the Laboratory-Activated Serratia Phage (LASφ) on the *S. symbiotica* CWBI-GFP ancestor. The lysate was made from a revived sample of the LT-02 endpoint population. Ten-fold serial dilutions of lysate, beginning on the left, were spotted on top agar containing host cells. This plate was incubated at ambient room temperature for 48 hours before imaging. (B) Representative TEM images of phage particles in lysate from the LT-02 endpoint population. Scale bars are 50 nm. (C) Coverage of DNA sequencing reads mapped to the CWBI-GFP genome from a pooled lysate of all 12 endpoint LT populations which contained LASφ phage. (D) Coverage of DNA sequencing reads mapped to the CWBI-GFP genome from a sample of the supernatant of ancestor cells treated with mitomycin C (MC).

To identify the reactivated phage and ascertain its origin, we pooled lysates made from all LT populations and extracted and sequenced their DNA. The CWBI-GFP genome contains 11 putative prophage regions, 10 of which are predicted to encode intact prophages (**Table 1**). Sequencing reads from the pooled phage lysates mapped to coordinates that matched a prediction of a 36.7-kb prophage at coordinate 1,323,010 (**Fig. 3C**). This genomic region is predicted to encode a complete copy of a P2-like prophage in the family Peduoviridae. We named this phage, which recurrently became lytic during our evolution experiments at the lower temperature, the Laboratory-Activated Serratia Phage (LASφ).

**Table 1.** Predicted prophage regions in the chromosome of the *S. symbiotica* ancestor.

| Designation | Position | Size (kb) | Proteins | Completeness | GC% | Annotation |
| --- | --- | --- | --- | --- | --- | --- |
|  | 28900 | 41.7 | 59 | Intact | 52.46 | PHAGE_entero_phiT5282H_NC_049429 |
|  | 221511 | 34.9 | 30 | Questionable | 53.05 | PHAGE_Escher_RCS47_NC_042128 |
|  | 441049 | 35.8 | 50 | Intact | 55.31 | PHAGE_Haemop_SuMu_NC_019455 |
| MC | 771733 | 48 | 78 | Intact | 49.34 | PHAGE_Salmon_118970_sal3_NC_031940 |
|  | 914879 | 35.7 | 52 | Intact | 55.89 | PHAGE_Haemop_SuMu_NC_019455 |
| LASϕ | 1323010 | 36.7 | 45 | Intact | 52.29 | PHAGE_Salmon_Fels_2_NC_010463 |
|  | 1884586 | 18.4 | 24 | Intact | 47.20 | PHAGE_Enterо_P1_NC_005856 |
|  | 1921934 | 89.6 | 102 | Intact | 50.27 | PHAGE_Salmon_118970_sal3_NC_031940 |
|  | 2111306 | 44.2 | 43 | Intact | 52.79 | PHAGE_Enterо_phiT5282H_NC_049429 |
|  | 2236529 | 16.8 | 29 | Intact | 54.87 | PHAGE_Ralsto_RSA1_NC_009382 |
|  | 2465690 | 17.8 | 21 | intact | 49.05 | PHAGE_Mannhe_vB_MhS_535AP2_NC_028853 |

LASϕ is the prophage region with nearly all DNA sequencing coverage in a pooled phage lysate from evolved samples. MC is the prophage region with DNA sequencing coverage in the supernatant of cultures of the ancestor treated with mitomycin C. Positions are for the chromosome of the *S. symbiotica* CWBI-GFP genome assembled in this study (**Data File S1**).

Next, we tested whether LASφ replication was activated by DNA damage by sequencing DNA from the supernatant of a CWBI-GFP culture exposed to mitomycin C. DNA in this sample was enriched in sequences that mapped to the approximate coordinates 772,000–819,000 of the CWBI-GFP genome (**Fig. 3D**). This region is predicted to encode an intact prophage in family *Myoviridae* related to bacteriophage 118970_sal3 (**Table 1**) (33). Therefore, we conclude that LASφ is not activated by the SOS response and that the CWBI genome encodes at least this one other intact prophage that is capable of entering the lytic cycle.

We surveyed the genomes of eleven other *S. symbiotica* strains to see if they harbored prophages related to LASφ. These strains exhibit a range of associations with aphids. Some are free-living gut pathogens like CWBI, and others are host-restricted facultative or co-obligate endosymbionts. There was at least one prediction of a prophage in the genome of every strain (**Table S1**). As expected for host-restricted genomes that have undergone reductive evolution, the aphid endosymbiont strains (AURT-53S, PCOLA-89S, IS, Tucson, PLYR-94S, PERIS-14S, SCt-VLC, MCAR-56S) generally harbor fewer putative prophages than the free-living strains (CWBI, HB1, 24_1, apa8a1). There are an average of 4.1 in the former (range: 1–8) and 7.8 in the latter (range: 4–11), though this difference is not significant (*p* = 0.10, two-tailed Mann-Whitney U test). Prophages assigned to the *Peduoviridae* family are widespread in these genomes (22 of 64 total prophages), but only the genomes of the culturable HB1 and 24-1 strains have prophages that appear to be closely related to LASφ from CWBI.

### Evolved LT clones are resistant to LASφ

Given the cell death and growth delays we observed in the LT populations, we expected there to be strong selection for the evolution of resistance to LASφ. To test whether this was the case, we made phage lysate from the 15-passage sample of population LT-02. We tested the effect of adding this lysate or the same lysate after it was heated to 90°C, on the growth of cultures of the ancestor and the LT clones (**Fig. 4**). The ancestor showed a normal growth curve when exposed to the heat-inactivated lysate but did not develop turbidity when subjected to phage lysate. By contrast, exposing most evolved clones to active phage lysate caused very little difference in their growth. The exception was the LT-07 clone, which was also inhibited by the phage. In conclusion, all but one of the evolved LT clones evolved resistance to LASφ.

**Fig. 4.**
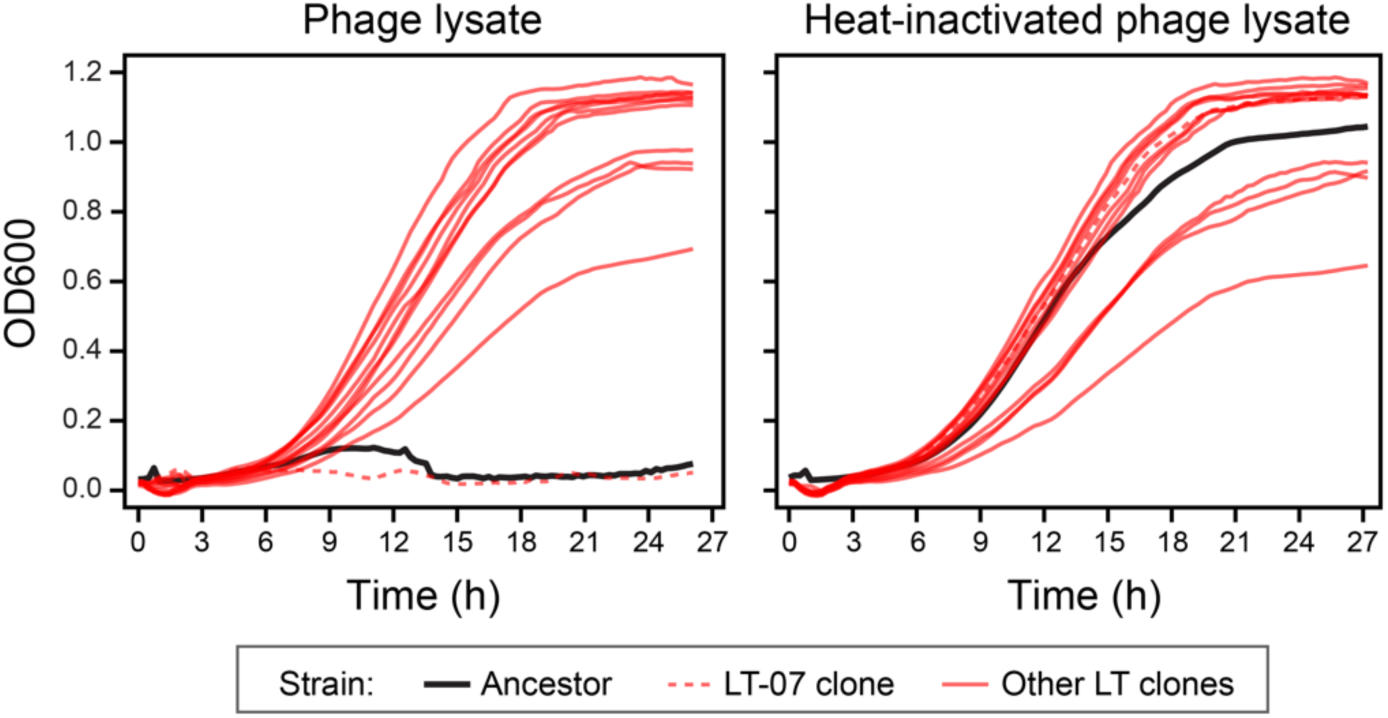
Evolved clones are resistant to LASφ. Growth curves were recorded as mean optical density at 600 µm (OD600) versus time in a microplate reader. Averages for replicate wells of clones (*n* = 3) and ancestor (*n* = 9) are shown. Standard errors on OD600 measurements were < 0.05 for all clones at all time points. Treatment with phage lysate from a sample of the endpoint LT-02 population inhibits growth of the CWBI-GFP ancestor and the LT-07 clone but not any of the other characterized LT clones. Inactivating the phage lysate by heating it abolishes this inhibition.

### Genome evolution

Next, we sequenced the genomes of all LT and HT endpoint clones to understand how they adapted to the laboratory environment and determine the genetic basis of resistance to LASφ. There were 1.1 and 0.8 mutations on average in each LT and HT clone, respectively (**Table 2**). The genomes of most clones had exactly one mutation, except two had two mutations each (LT-05 and HT-11) and three—all in the high-temperature treatment—had no mutations (HT-01, HT-02, and HT-08). Base substitutions were the most abundant type of mutation, with 12 total across both temperature conditions, accounting for 57% of the 21 total mutations. All base substitutions were nonsynonymous mutations in protein-coding genes. There were also four small insertions and deletions of ≤ 50-bp, two of which were in homopolymer repeats. Lastly, we observed three IS element insertions, a 54-bp deletion, and a 27.4-kb deletion.

**Table 2.**
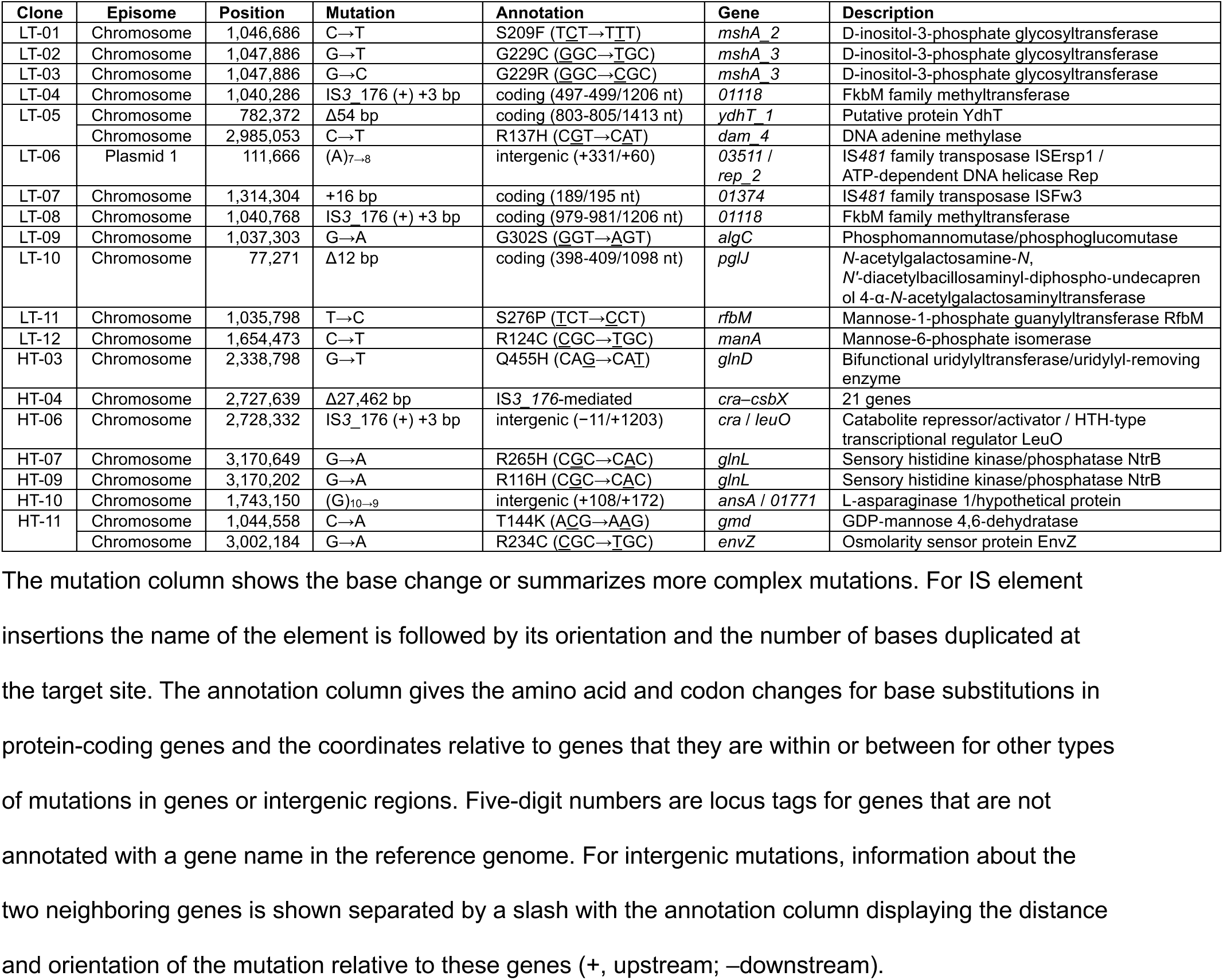
Mutations in evolved clonal isolates.

### Mutations affecting LPS biosynthesis are associated with phage resistance

Most mutations in the LT evolved clones appear to be associated with synthesis or modification of lipopolysaccharide (LPS) or other extracellular polysaccharides (**Table 2**) (34–39). LT-09, LT-11, and LT-12 had mutations in genes that are involved in the biosynthesis of GDP-mannose, a key LPS precursor (34, 37–39). LT-01, LT-02, LT-03, and LT-10 had mutations in glycosyltransferases that could link monosaccharide units or attach them to lipids (35, 36). LT-04 and LT-08 had mutations in the same FkbM family methyltransferase that could modify sugars displayed on the cell surface (40). As these were often the sole mutations in these clones and they are broadly convergent in terms of their possible functions, it seems likely that these mutations confer resistance to LASφ by altering polysaccharides on the bacterial cell surface. Clone LT-07 was the only endpoint clone at RT that was not resistant to LASφ (**Fig. 4**), and it did not have mutations in any of these pathways.

Clone HT-11, which exhibited the settling phenotype observed in this population, had two mutations. One was in a mannose biosynthesis gene *gmd*, and one was in the osmolarity sensor *envZ* (41, 42). It is possible that stochastic reactivation and the evolution of resistance to LASφ also caused the evolution of the unique phenotype displayed by this population, through the way in which it altered its cell surface. None of the other evolved HT clones had mutations in genes with a direct connection to polysaccharide biosynthesis or modification. Instead, nearly all mutations in HT clones are in genes involved in regulation of nitrogen metabolism or central carbon metabolism, including convergent mutations affecting *glnL* and *cra* (43–48).

### IS*3* elements mediated gene loss in CWBI evolved at both temperatures

The CWBI genome is predicted to have 232 insertion sequence (IS) elements from fifteen families, though 124 of these (53.8%) appear to be incomplete or have inactivating mutations. The most frequent families are IS*3* (100 total, 49 complete), IS*256* (37 total, 12 complete), and IS*4* (23 total, 21 complete). In endpoint clones from the evolution experiment, we only observed activity of one specific IS*3* subfamily with 48 copies of which 40 appear to be active. We observed a large deletion that led to the loss of 21 genes in HT-04 mediated by insertion of a new copy of this IS element in the *csrA* gene followed by recombination with a preexisting copy overlapping the end of the *csbX* gene. The same IS*3* element inserted into genes in LT-04 and LT-08 and into an intergenic region in HT-06. No other IS element families mediated deletions or were observed to insert new copies during the evolution experiment.

### Evolved LT clones are less virulent than the ancestor when injected into aphids

To determine if mutations that accumulated during the evolution experiment attenuated CWBI virulence, we injected cohorts of 4th instar aphids with each of the evolved LT clones and compared their survival to aphids injected with the ancestor strain. Under these conditions aphids rapidly become systemically infected with the *S. symbiotica* CBWI-GFP ancestor (**Fig. 5A**). Whereas a majority of aphids survive for ∼12 days after mock injections with buffer (**Fig. 5B**), most die within 3-4 days after injection with the CWBI-GFP ancestor (**Fig. 5C**).

**Fig. 5.**
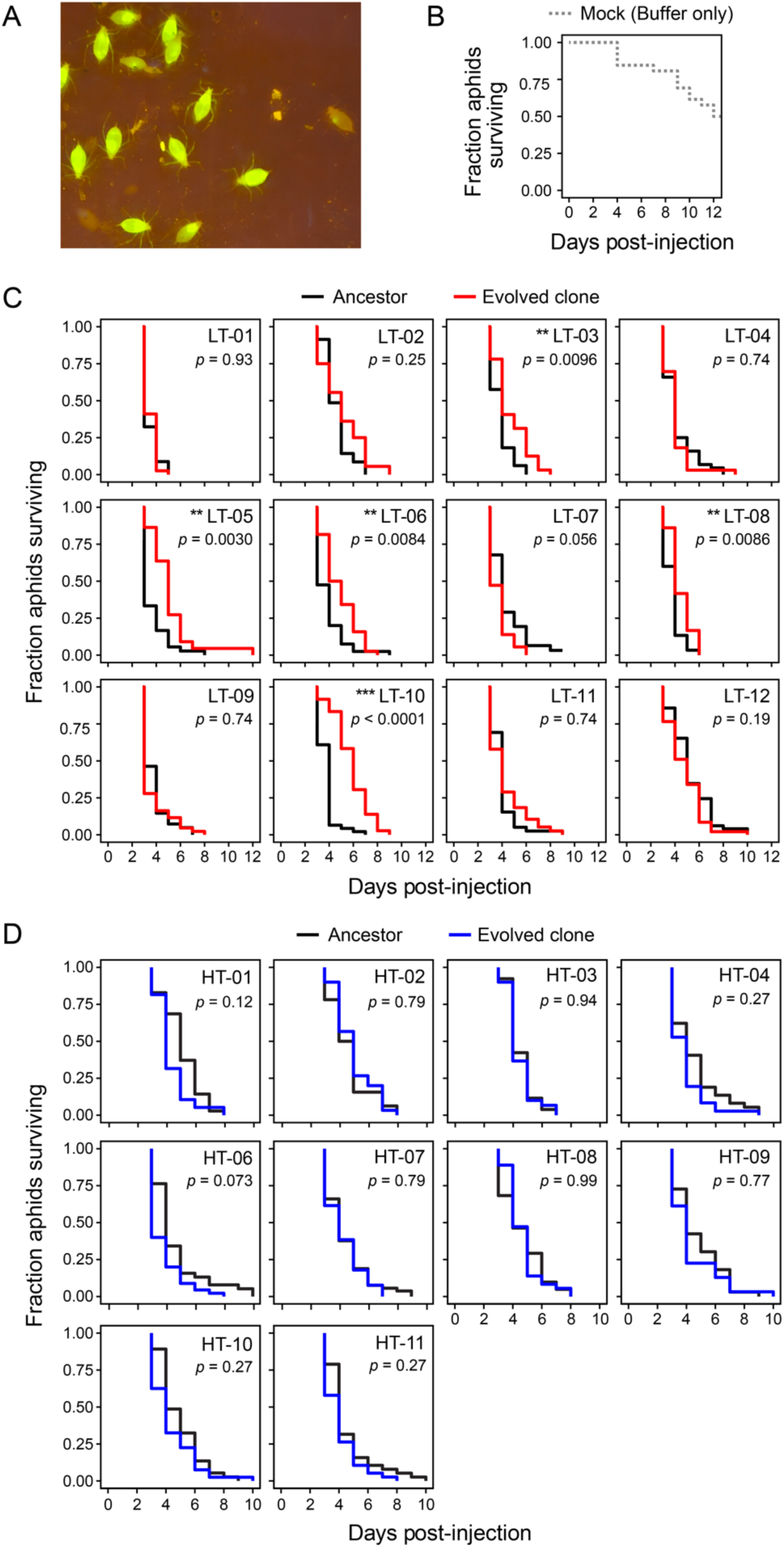
Some LT-evolved clones exhibit slower killing of aphids. (**A**) The CWBI-GFP ancestor rapidly and systemically infects when ∼10^4^ bacterial cells are microinjected into aphids. GFP fluorescence in living aphids was imaged on a blue-light transilluminator with an orange filter at 4 days post-injection. (**B**) Survival curve for a cohort of 32 fourth-instar aphids mock injected with buffer. (**C**) Survival curves for cohorts of 40-50 fourth-instar aphids injected with ∼10^4^ *S. symbiotica* cells of clones beginning at 2 days post-injection. The 12 endpoint clones from the LT populations were each tested at the same time as a paired ancestor control. \**p* < 0.05, \*\**p* < 0.01, \*\*\**p* < 0.001 for two-tailed log-rank tests comparing Kaplan-Meier survival curves for each clone and its paired ancestor control after adjusting p-values for multiple comparisons using the Benjamini-Hochberg procedure. (**D**) Survival curves for the 10 endpoint clones from the HT populations and their paired ancestor controls. Details are as in **C**.

We observed significantly longer aphid survival after injecting 5 of the 12 LT clones evolved at ambient room temperature compared to paired injections with the ancestor strain (*p* < 0.05, two-tailed log-rank tests with Benjamini-Hochberg correction) (**Fig. 5C**). The most dramatic changes were for LT-10 and LT-05. These clones killed aphids an average of 2.19 and 1.48 days more slowly than the ancestor, respectively. The evolved clone that was still susceptible to phage, LT-07, was the only one that exhibited the opposite behavior. Aphids died more quickly by an average of 0.55 days after they were injected with this clone, though this difference was only marginally statistically significant (*p* = 0.056), By contrast, none of the aphids injected with the HT clones evolved at 30°C survived significantly longer that aphids injected with the ancestor (**Fig 5D**). Two clones, HT-01 and HT-06, killed aphids more quickly, by an average of 0.72 and 0.85 days faster than the ancestor, respectively. This increased virulence was also only marginally significant at best (*p* = 0.12 and *p* = 0.073, respectively),

A faster growth rate or higher final cell density for a given evolved strain when cultured *in vitro* might generally lead to trade-offs that make it less virulent to aphids (49–51). Alternatively, improved growth in the rich media used in the evolution experiment might align with improved *in vivo* growth within an aphid’s body and therefore increase virulence (49–52). We did not find support for either of these possibilities. There was no significant correlation between the *in vitro* growth rates or carrying capacities fit from growth curves and aphid survival following injections for either the LT- or the HT-evolved clones (*p* > 0.05, *t*-tests on slope terms in linear regressions) (**Fig. S1**). We also could not discern a pattern with respect to the presence of mutations in certain genes or pathways in LT-evolved clones that exhibited significantly reduced virulence versus the others. Finally, we did not observe vertical transmission of ancestor or evolved *S. symbiotica* to aphid offspring in these experiments, which is in line with the delay of 10 days post-injection before infected offspring are observed in prior experiments with *S. symbiotica* (10). Nevertheless, the substantial changes in survival for certain clones indicate that evolution in this short laboratory experiment was able to substantially attenuate the virulence of this *S. symbiotica* protosymbiont in some cases.

### Evolved attenuated strains are still transmitted to aphid embryos

To test whether mutations in LT-evolved strains, including some with significantly reduced virulence, prevented *S. symbiotica* from being transovarially transmitted to embryos, we performed a new set of injection experiments. This time we injected with fewer bacterial cells (∼10^3^) and left these aphids undisturbed on larger plants to maximize their lifespans. On the seventh day post-injection, we used confocal microscopy to look for fluorescent bacterial cells in the developing aphid embryos. The ancestor and all three evolved clones we tested (LT-08, LT- 09, and LT-10) were capable of colonizing embryos (**Fig. 6**). Overall, there seemed to be fewer LT-08 and LT-10 (**Fig. 6C,E**) bacterial cells per embryo compared to cells of the ancestor and LT-09 (**Fig. 6B,D**), which tracks with the relative virulence of these strains (**Fig. 5**).

**Fig. 6.**
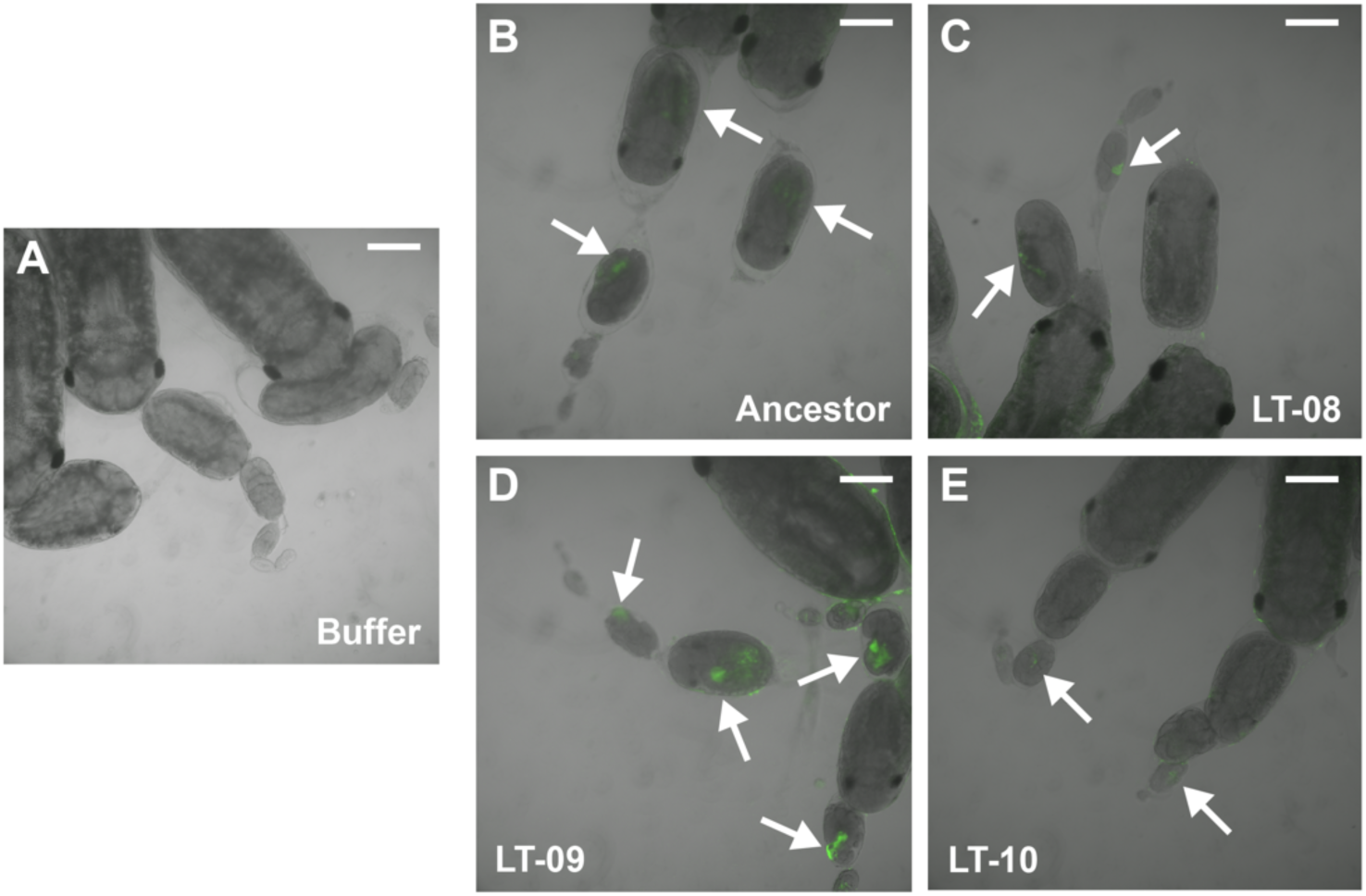
Attenuated evolved clones are still transovarially transmitted to aphid embryos. Embryos in ovarioles dissected from adult aphids seven days after injecting them as fourth instars with (A) buffer (mock injection) or ∼10^3^ cells of the (B) CWBI-GFP ancestor, (C) evolved clone LT-08, (D) evolved clone LT-09, or (E) evolved clone LT-10. Arrows point out fluorescent patches within embryos typical of transovarial transmission of GFP-tagged *S. symbiotica* cells. Scale bars are 200 µm.

## DISCUSSION

Our evolution experiments with culturable *S. symbiotica* strains began with two major goals related to investigating routes by which this protosymbiont strain could mutate to become a stably vertically inherited endosymbiont of aphids. First, we wanted to understand whether adaptation to a nutrient-rich laboratory environment will attenuate the growth and virulence of CWBI when it is injected into aphids. Second, we wanted to characterize the activities of different families of mobile elements in the CWBI genome to understand their relative contributions to inactivating and deleting genes, which could also attenuate CWBI virulence. In general, one can imagine distinct routes by which mutations resulting from either non-adaptive genetic drift or adaptive decay due to trade-offs between growth in different environments transform a pathogen into an endosymbiont (17, 18, 20, 21, 31). Recapitulating this process could inform our understanding of how and when each pathway contributes to this transition.

Insertion sequence (IS) elements are simple bacterial transposons that can inactivate genes and mediate large chromosomal rearrangements (53, 54). Given the prevalence and diversity of IS elements in the *S. symbiotica* CWBI genome, we expected that IS-mediated mutations would dominate during our evolution experiment. However, only one subfamily of IS*3* elements was active. Of 21 total mutations, just three were new insertions of this transposon and one was a 27-kb deletion mediated by this transposon. The IS-mediated fraction of total mutations in our experiment with CWBI (19%) is not as high as that seen in evolution experiments with laboratory strains of *E. coli* (30%) (55) and *Acinetobacter baylyi* (41%) (56), even though these bacteria have fewer transposon copies than CWBI in their genomes. Thus, perhaps surprisingly, the genome of CWBI does not appear to be unusually unstable. Nevertheless, we would expect IS elements—both existing copies and new ones—to have an outsize role in catalyzing large-scale deletions if our evolution experiment continued.

Recently, within-host evolution experiments showed that a single mutation in the *crp* master regulator of central carbon metabolism can make *E. coli* an effective substitute for the native *Pantoea* gut symbiont of the stinkbug *Plautia stali* in supporting its nutrition and development (57). Similar global regulatory mutations affecting different growth switches, particularly stationary phase to exponential phase transitions and biofilm versus planktonic lifestyles, could greatly alter CWBI interactions with aphid hosts. In this vein, it is interesting that we observed a mutation in the upstream region of *cra*, another master regulator of central carbon metabolism, and an IS-mediated deletion overlapping this gene in two of the evolved clones from the higher-temperature treatment. This condition also led to the evolution of *glnD* and *glnL* mutations that are expected to affect global regulation of nitrogen metabolism. How these mutations affect *S. symbiotica* growth in aphids and host survival remains to be tested.

Emergence of the Laboratory-Activated *Serratia* Phage (LASφ) drove the initial stages of our evolution experiment in the lower temperature (LT) treatment. Typically, prophage activation occurs when bacteria experience stress, whether from nutrient deprivation or DNA damage (58–60). Temperature can be one such stress, though classical *E. coli* prophage models are based on artificial temperature-sensitive repressor mutants that activate at high temperature (e.g., λ cI857) (61). A study that examined and evolved *Serratia marcescens* (ATCC 13880) at different temperatures in the laboratory (38°C, 31°C, and 24-38°C fluctuating) also found increased activation of a P2-like prophage associated with lower temperatures (62), suggesting this response may be conserved among this class of *Serratia* phages.

Reactivation of LASφ created a strong selective pressure for CWBI to evolve resistance in our LT populations, which occurred through mutations affecting lipopolysaccharide (LPS) biosynthesis. The *S. marcescens* laboratory evolution experiments also resulted in mutations in LPS biosynthesis, though their effects on phage resistance were not tested (62). Mutations in other bacterial pathogens that confer resistance to phage often affect similar pathways and result in trade-offs that reduce fitness *in vitro* and *in vivo* when phage is not present (63, 64). For example, *Listeria monocytogenes* that evolved mutations in teichoic acid glycosylation genes that conferred resistance to a phage was less virulent in mice and exhibited a reduced ability to invade cells (65). Similar trade-offs have been observed in phage-resistant mutants of *Vibrio cholerae* (66), *Shigella flexneri* (67), and *Staphylococcus aureus* (68), among others. Some of the clonal endpoint isolates from our experiment that were resistant to LASφ grew more slowly than the ancestor, suggesting that they exhibit fitness trade-offs for phage resistance *in vitro*, but not all did. The nutrient-replete laboratory conditions of our evolution experiment likely allowed phage resistance to evolve via mutational routes that would not be favored in complex natural environments in which bacteria experience other stresses and selection pressures.

Prophages have been found to have important roles in other insect endosymbionts, but typically this occurs when the functions of prophage genes are co-opted to benefit the symbiont and/or its host. The APSE prophage in the genome of the endosymbiont *Hamiltonella defensa* encodes a toxin that has been shown to protect aphids from parasitoid wasps, thereby contributing to persistence of this facultative symbiont in some aphid populations (69–71). Genes expressed from WO prophage regions in the genomes of *Wolbachia* mediate how this endosymbiont manipulates reproduction in other types of insects (72, 73).

We found that some of our LT-evolved *S. symbiotica* clones with LASφ-resistance mutations were less virulent when injected into aphids. Similarly isolates from populations of the *S. marcescens* evolution experiment that encountered the lower temperature had reduced virulence when injected into insects (*Galleria mellonella* larvae) (62). Overall, these results suggest additional dimensions to how bacteria-phage coevolution might shape the emergence of new endosymbionts. Culturable *S. symbiotica* that are naturally acquired by aphids from their diet sometimes escape the gut and systemically infect their bodies (8). Prophage activation, phage predation, or the pleiotropic effects of phage-resistance mutations may sometimes reduce the titer or growth rates of these bacteria, thereby increasing the chances that they are able to evolve and establish as stably vertically inherited symbionts before they kill their hosts. It is also possible that mutations that confer phage resistance could prevent the establishment of endosymbionts. For instance, changes in LPS or loss of cell-surface proteins that act as phage receptors in resistant mutants could affect interactions with aphid embryos that lead to endocytosis of some but not other bacteria (10, 74). We did not observe this potential trade-off: phage-resistant LT-evolved clones that were less virulent were still able to colonize aphid embryos. Additional evolution or genetic engineering will be needed to understand whether any of the mutations we observed are potential steppingstones to new aphid endosymbionts.

## MATERIALS AND METHODS

### Culture conditions

We used CWBI-GFP, a strain of *S. symbiotica* CWBI-2.3^T^ with genes for GFP and kanamycin resistance integrated into its chromosome (10). CWBI-GFP was cultured in trypticase soy broth (TSB) or on trypticase soy agar (TSA) (BD Biosciences) with 50 µg/mL kanamycin (Kan).

Unless specified otherwise, cultures were grown at in an incubator at 30°C or at the ambient temperature in our laboratory (22-25°C) with orbital shaking at 200 r.p.m. over a one-inch diameter in 5 mL of TSB in 150 × 18 mm test tubes. Samples were cryopreserved by adding glycerol to a final concentration of 16% (v/v) and stored at −80°C.

### Evolution experiment

Twenty-four colonies of CWBI-GFP from a TSA+Kan plate were used to inoculate separate cultures of TSB+Kan to begin the separate populations of the evolution experiment. Half were grown at ambient temperature (LT populations). Half were grown at 30°C (HT populations).

Upon achieving saturated growth, as judged by visual inspection, 5 µL of each culture was transferred to a new test tube containing fresh TSB+Kan. LT and HT populations were passaged 15 and 30 times, respectively. Samples of these cultures were cryopreserved every five passages. To isolate endpoint clones, glycerol stocks of the final populations were struck out on TSA+Kan plates. Then, single large colonies were picked from each plate into TSB+Kan, and these cultures were grown to saturation and cryopreserved.

### Growth curves

Cultures of evolved clones or populations were revived from cryopreserved samples and grown to saturation were diluted 1:100 into TSB+Kan. From each diluted culture, 300 µL was pipetted into a clear, flat-bottomed 96-well plate (Corning). The final layout of the plate included three replicates of a TSB+Kan blank, nine replicates of the ancestor, and three replicates of each evolved clone. Growth curves were measured in a Tecan Infinite M200 PRO plate reader.

Measurements of optical density at 600 nm (OD600) were recorded every 15 minutes with 14 minutes of orbital shaking at 200 r.p.m. between each reading for 48 hours. The temperature control in the instrument was set to 23⁰C for LT samples and 30⁰C for HT samples, but actual temperatures varied to as high as 27⁰C or 32⁰C, respectively, during runs due to heating from operation of the instrument. Growth curves were performed separately for each set of populations and clones evolved at different temperatures. Statistical analysis and plotting were performed in R using the *tidyverse, ggplot2*, *statix*, and *growthcurver* packages (75).

### Phage lysate preparations

For each lysate, 5-25 mL of saturated culture was mixed with an equal volume of chloroform in a centrifuge tube. After inverting vigorously for 15 seconds to mix, tubes were centrifuged for 5 minutes at 3,220 × g at ambient temperature. The supernatant was transferred to a fresh tube and 100 µL of chloroform mixed in by inverting the tube. The resulting lysate was stored at 4°C.

### Phage plaque assay

CWBI-GFP was grown to an OD600 of ∼0.4 in TSB+Kan at ambient temperature. Phage lysates were serially diluted 10-fold using TSB in a 96-well plate up to a maximum dilution of 10^5^. TSA top agar was made by halving the concentration of agar used to make TSA base media. Top agar was melted, then 25 mL was poured into a 100 mL flask and left to cool until warm.

Kanamycin was added along with 1.5 mL of culture. The mix was swirled briefly to mix, then poured onto the top of a TSA+Kan plate. Upon cooling, 2 µL of the lysate dilutions were pipetted onto the cooled top agar. After allowing the spots to dry, the plates were incubated at room temperature. Phage plaques became visible after 24 h and were more pronounced after 48 h.

### Transmission electron microscopy imaging

To isolate phage particles, 4 mL of lysate from population LT-02 was mixed with 1 mL of phage precipitation solution (20% w/v PEG 8000, 2.5 M NaCl) and mixed by inverting the tube for 15 seconds. The solution was stored at 4°C overnight then centrifuged for 30 minutes at 10,000 × g at 4°C. Supernatant was removed, and the phage pellet was resuspended and diluted in 1× phage resuspension buffer (1M NaCl, 10 mM Tris•HCl pH 7.5, 0.1 mM EDTA). The phage suspension was added to a TEM grid, washed with water, and stained with 2% uranyl acetate. Images were obtained using a FEI Technai transmission electron microscope at the University of Texas at Austin Center for Biomedical Research Support (CBRS) microscopy facility.

### Mitomycin C treatment

The CWBI ancestor strain was grown in TSB+Kan at room temperature to saturation from a cryopreserved stock. In a 50 mL Erlenmeyer flask, 110 µL of saturated culture was added to 11 mL of fresh TSB+Kan. Using a Genesys 150 UV-Vis spectrophotometer (ThermoFisher Scientific), initial OD600 values were measured using 1 mL of the culture. After 3 hours at room temperature under standard shaking conditions, the culture was split by pipetting 5 mL into two test tubes. In one test tube, 5 μL of a 0.6 mM solution of mitomycin C (MilliporeSigma) dissolved in water was added to one of the tubes (0.6 µM final concentration). After 24 hours of additional growth, 1 mL of the culture treated with mitomycin C was removed and centrifuged for 10 minutes at 4,000 × g at 4⁰C. From the supernatant, 857 µL was removed, added to 6 mL of DNA binding buffer from the DNA Clean & Concentrator kit (Zymo Research). DNA purified with this kit and eluted in 25 µL of DNAse/RNAse free water. The DNA concentration was measured using an Invitrogen Qubit 4 Fluorometer (ThermoFisher Scientific).

### Phage susceptibility testing

Phage lysate was prepared from 25 mL of culture as described above. Approximately 2 mL of lysate was inactivated by heating to 90°C for 10 minutes. For each growth curve replicate, 100 µL of lysate was added to 400 µL of TSB+Kan in a 1.7 mL Eppendorf tube, and then 5 µL of saturated ancestor strain was added to the tube. After mixing by inverting, 300 µL from the tube was added to a single well on a 96-well plate. Nine replicate wells of ancestor and three replicates well of each evolved clone were each tested both with lysate and with heated lysate. Nine blank wells of TSK+Kan were included. Growth curves were carried out and analyzed as described above.

### Phage DNA purification

Isolation and precipitation of phage particles was performed as described for TEM imaging. To eliminate carryover of nucleic acids from host cells, 40 µL of 10× DNase I buffer (New England Biolabs) was added to 360 µL of phage sample in resuspension buffer. Then, 1 μL DNase I (2000 U/mL, New England Biolabs) and 1 μL RNase A (20 mg/mL, Invitrogen PureLink™ RNase A) were added. Following incubation at 37°C for 30 min, 10 μL of 0.5 M EDTA (pH 8.0) was added to halt digestion. To remove enzymes and purify DNA from phage particles, the digested sample was phenol extracted and ethanol precipitated. The final DNA pellet was resuspended in 10 mM Tris•Cl (pH 7.5) and stored at −20°C.

### Genome sequencing

TSB+Kan cultures of the ancestor strain and evolved clones were grown to saturation at their respective temperatures. For Illumina sequencing, genomic DNA (gDNA) was isolated from 1 mL of culture using a PureLink Genomic DNA Mini Kit (Invitrogen). A Qubit 2.0 Fluorometer with the broad range assay was used to measure DNA concentrations. Up to 10 ng of gDNA was used as input to the IDT xGen DNA Lib Prep EZ using the xGen Deceleration Module to isolate larger fragment sizes according to the manufacturer’s instructions except that all reactions took place at 20% reaction volume. Libraries were sequenced in paired-end 150-bp mode using unique 6-bp dual index sample barcodes. Bacterial genomes were sequenced on a HiSeq X Ten at Psomagen (Rockville, MD). Phage genomes were sequenced on an iSeq 100.

For nanopore sequencing, high molecular weight genomic DNA was isolated using the 20/G Genomic-tips (Qiagen) or the Quick-DNA HMW MagBead Kit (Zymo research). For each sample, 325-400 ng of this gDNA was prepared for sequencing using a Rapid Barcoding kit (Oxford Nanopore Technologies). Samples were sequenced on a MinION Mk1C using V9.4.1 flow cells. Onboard basecalling using Guppy (V 5.0.13) was done in fast mode.

### Ancestor genome assembly and annotation

CWBI-GFP nanopore reads were trimmed of adaptors using Porechop and filtered based on quality and length using Filtlong (76). Illumina reads were trimmed using fastp (77). We used Trycycler (78) to generate a consensus assembly from Flye (79), Miniasm+Minipolish (80), Canu (81), NECAT (82), Raven (83), and Unicycler (84) assemblies of long read subsets. The final assembly was iteratively polished, first using Medaka (Oxford Nanopore Technologies) with nanopore reads and then using Polypolish (85), POLCA (86), and *breseq* (87) with Illumina reads. Finally, we used Prokka (88) and ISEscan (89) to annotate genes and IS elements in the final genome assembly (**Data File S1**). Prophages were predicted using the PHASTEST webserver (v3.0) (90–92) and assigned to taxonomic groups by running PhaGCN (93) on the PhaBOX webserver (v2.0) (94).

### Read mapping and mutation calling

We predicted mutations in evolved clones relative to the annotated CWBI-GFP genome using *breseq* (v0.38.1) (87). This pipeline uses bowtie2 (95) for read alignment. We also mapped reads from phage and mitomycin C samples to the reference genome and examined read coverage using *breseq*. In these samples, some regions of the LASφ prophage and the prophage activated by mitomycin C share short stretches of exact homology with other prophage regions that initially resulted in ambiguous mapping of some reads. To create the final coverage graphs, we used an artificial reference sequence that avoided these artifacts by including just the first half of the genome. Coverage plots zoomed to the prophage regions were created using the bam2cov post-processing command from *breseq*.

### Aphid injection

Ancestor and evolved clones were inoculated into TSB+Kan from glycerol stocks and grown at ambient temperature. Cells were pelleted by centrifugation at 6800 × g for 3 min and then washed three times in 1× PBS with 1 min of centrifugation each time. Cell pellets were resuspended in buffer A (25 mM KCl, 10 mM MgCl_2_, 250 mM sucrose, 35 mM Tris-HCl, pH 7.5) at an OD600 of 1.0. For injection, 40-50 fourth-instar aphids per treatment were chilled overnight at 4⁰C, then injected at the base of the hindleg with <0.1 µL of resuspended cells using pulled 5 µL calibrated micropipettes (Drummond) loaded into a IM-400 electric microinjector (Narishige) (MPa = 0.037, 0.2 seconds). To estimate the number of cells injected into each aphid, the microinjector was used to dispense <0.1 µL of resuspended cells into tubes containing 50 µL of buffer A, 1:10 serial dilutions were spot-plated, and CFUs were counted. The number of cells injected per aphid was approximately ∼10^4^ cells. Immediately post-injection, aphids were placed onto 5- to 5.5-inch plants and allowed to recover for 24 h. Counting of surviving aphids began 48 hours post-injection and continued daily. For counting, aphids were gently removed from the plants and placed under a blue-light illuminator to also monitor them for colonization. Aphids were placed back on plants immediately after counting. Statistical tests and plotting of survival curves were conducted in R using the *survminer* and *survival* packages (96).

### Confocal microscopy

Fourth instar (7-day old) aphids were injected with buffer A, the ancestor strain, clone LT-08, clone LT-09, or clone LT-10 (30 aphids per treatment, ∼1000 cells injected per aphid) and immediately left to feed on large plants (>12 inches), undisturbed. After seven days, the grown-up adult aphids were removed, checked for fluorescence, and dissected in 1× PBS. Ovarioles were mounted in 60 µL 1×PBS on concave glass slides (ZTBH #7103) and imaged using a Nikon AXR upright confocal microscope. Images were taken at 10× and 20× magnification using GFP settings.

## DATA AVAILABILITY

Whole-genome sequencing reads have been deposited in NCBI Sequence Read Archive (PRJNA1111250).

## ACKNOWLEDGMENTS

We thank Daniel Deatherage for assistance with DNA sequencing; Victor Li for assistance with phage experimental techniques; and Michelle Mikesh of the UT Microscopy Core for assistance with TEM sample preparation and operation.

