## Supplementary material for "Evolution in response to prophage activation attenuates the virulence of culturable *Serratia symbiotica* relatives of aphid endosymbionts": Figure S1

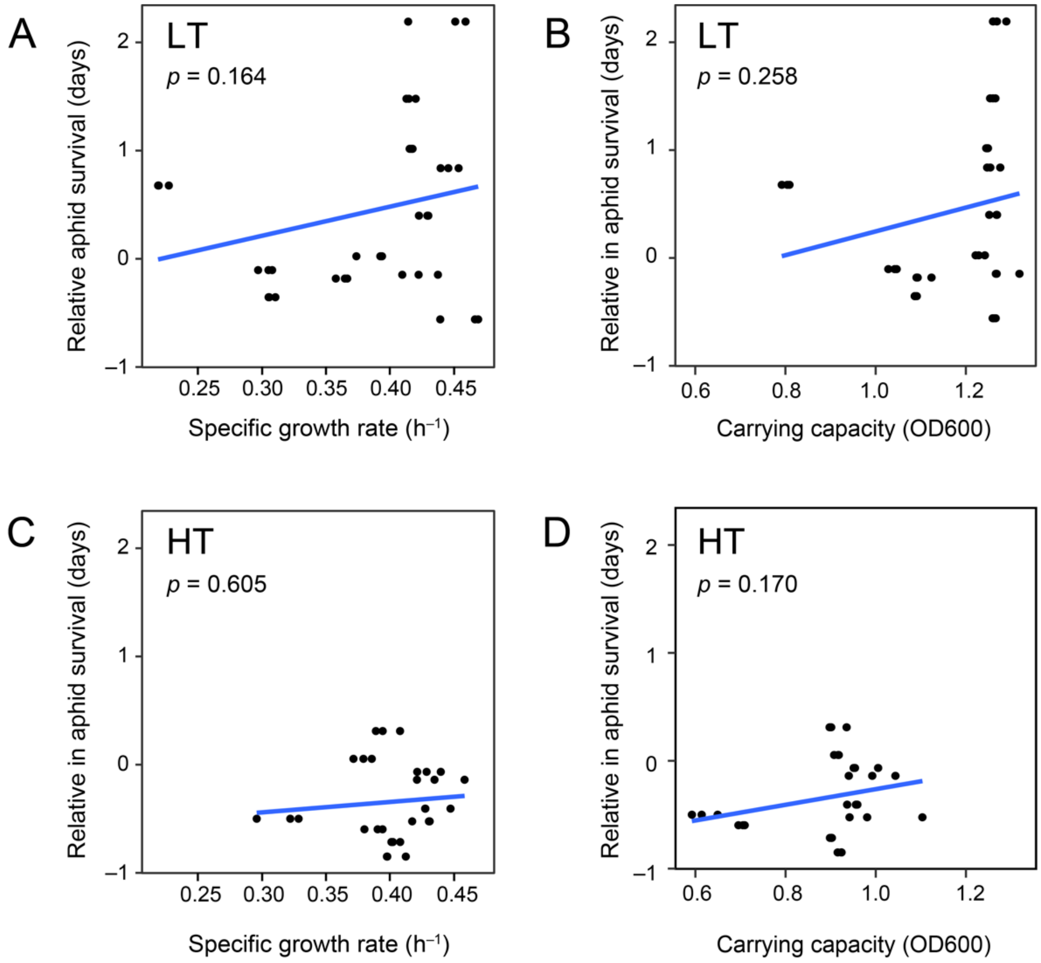


**Fig. S1. Growth rates and carrying capacities of evolved *S. symbiotica* clones cultured *in vitro* are uncorrelated with their virulence when injected into aphids.** Linear regressions were conducted between the average number of days aphids injected with the evolved clone survived relative to aphids injected with the ancestor strain (areas under the curves in **Fig. 5**) and parameters fit from each of three separate growth curve replicates for each clone. Each $p$-value is for rejecting the null hypothesis that there is no significant correlation ($t$-test that the slope of the fit line shown is not different from zero). (**A**) Correlation between growth rate and aphid survival for the LT-evolved clones. (**B**) Correlation between carrying capacity and aphid survival for the LT-evolved clones. (**C**) Correlation between growth rate and aphid survival for the HT-evolved clones. (**D**) Correlation between carrying capacity and aphid survival for the HT-evolved clones.
